# Motor Control of Distinct Layer 6 Corticothalamic Feedback Circuits

**DOI:** 10.1101/2024.04.22.590613

**Authors:** Luis E. Martinetti, Dawn M. Autio, Shane R. Crandall

## Abstract

Layer 6 corticothalamic (L6 CT) neurons provide massive input to the thalamus, and these feedback connections enable the cortex to influence its own sensory input by modulating thalamic excitability. However, the functional role(s) feedback serves during sensory processing is unclear. One hypothesis is that CT feedback is under the control of extra-sensory signals originating from higher-order cortical areas, yet we know nothing about the mechanisms of such control. It is also unclear whether such regulation is specific to CT neurons with distinct thalamic connectivity. Using mice (either sex) combined with *in vitro* electrophysiology techniques, optogenetics, and retrograde labeling, we describe studies of vibrissal primary motor cortex (vM1) influences on different CT neurons in the vibrissal primary somatosensory cortex (vS1) with distinct intrathalamic axonal projections. We found that vM1 inputs are highly selective, evoking stronger postsynaptic responses in Dual ventral posterior medial nucleus (VPm) and posterior medial nucleus (POm) projecting CT neurons located in lower L6a than VPm-only projecting CT cells in upper L6a. A targeted analysis of the specific cells and synapses involved revealed that the greater responsiveness of Dual CT neurons was due to their distinctive intrinsic membrane properties and synaptic mechanisms. These data demonstrate that vS1 has at least two discrete L6 CT subcircuits distinguished by their thalamic projection patterns, intrinsic physiology, and functional connectivity with vM1. Our results also provide insights into how a distinct CT subcircuit may serve specialized roles specific to contextual modulation of tactile-related sensory signals in the somatosensory thalamus during active vibrissa movements.

**SIGNIFICANCE STATEMENT:** Layer 6 corticothalamic (L6 CT) feedback circuits are ubiquitous across mammalian species and modalities, and their activities have a strong influence on thalamic excitability and information throughput to the neocortex. Despite clear evidence of CT effects on the thalamus, we know relatively little about how CT cells themselves are regulated. Our results show that input from the primary motor cortex strongly excites a subclass of CT neurons in the primary somatosensory cortex that innervate both core and higher-order somatosensory nuclei rather than those exclusively targeting core somatosensory thalamus. The cortico-cortico-thalamic pathway formed by these connections establishes a circuit-level substrate for supporting CT influence operating under the guidance of ongoing motor activities.

## INTRODUCTION

Nearly all sensory signals enter the neocortex through the thalamus, and the neocortex, in turn, sends a massive projection back to the thalamus. A remarkable feature of this reciprocal arrangement is that descending layer 6 corticothalamic (L6 CT) axons outnumber ascending thalamocortical (TC) axons by 10:1 (Sherman and Koch, 1986), and L6 CT axons produce 30– 44% of all synapses onto TC neurons (Liu et al., 1995; Erisir et al., 1997; Van Horn et al., 2000). Due to this anatomical organization, L6 CT neurons are strategically positioned to modulate thalamic processing and TC communication.

Although much work has focused on how the CT pathway modulates thalamic excitability and sensory responses (for review, see (Briggs and Usrey, 2008; Briggs, 2020)), much less is known about the inputs controlling CT cells themselves. This question is of particular interest given many CT cells are silent and unresponsive to sensory stimuli *in vivo* (Tsumoto and Suda, 1980; Swadlow and Hicks, 1996; Sirota et al., 2005; Kwegyir-Afful and Simons, 2009; Egger et al., 2020). A further complication is that L6 CT neurons are morphologically and physiologically diverse (Tsumoto and Suda, 1980; Bourassa et al., 1995; Briggs and Usrey, 2009; Briggs et al., 2016; Stoelzel et al., 2017; Chevee et al., 2018). Despite limited thalamic inputs (Crandall et al., 2017), several studies suggest that CT cells receive non-sensory inputs from other cortical regions (Zhang and Deschenes, 1998; Hooks et al., 2013; Velez-Fort et al., 2014; DeNardo et al., 2015). Determining the afferent inputs targeting CT cells and the cellular and synaptic mechanisms underlying their activation is an important step toward understanding when and why CT circuits operate.

In the rodent vibrissal system, one important pathway that provides input to the vibrissal primary somatosensory cortex (vS1) is the vibrissal primary motor cortex (vM1) (White and DeAmicis, 1977; Porter and White, 1983; Cauller et al., 1998; Petreanu et al., 2009). vM1 projections terminate densely in infragranular layers of vS1 (Miyashita et al., 1994; Veinante and Deschenes, 2003), where they facilitate the neuronal responses of infragranular neurons *in vivo*, including L6 CT cells (Lee et al., 2008). Since vM1 activity correlates with vibrissa movement (Carvell et al., 1996; Hill et al., 2011; Sreenivasan et al., 2016), vM1 projections might function to communicate the motor plan to vS1 (Kleinfeld et al., 2006). Thus, L6 CT cells appear capable of modulating thalamic throughput under the guidance of vM1 during active vibrissa movements.

However, an understanding of how vM1 input facilitates CT activity is currently lacking. For example, CT neurons projecting to the ventral posterior medial nucleus (VPm), the core vibrissal division of the somatosensory thalamus, receive weak vM1 input compared to non-CT excitatory cells (Kinnischtzke et al., 2016). In theory, vM1 could indirectly excite CT cells by local circuit interactions (Zagha et al., 2013), but they receive relatively few local inputs (Thomson, 2010; Velez-Fort et al., 2014; Crandall et al., 2017). Alternatively, vM1 input could be biased towards specific CT neurons. In rodent vS1, L6 CT cells fall into two broad subclasses based on their intrathalamic axonal projections and position within L6 (Bourassa et al., 1995; Killackey and Sherman, 2003; Chevee et al., 2018). Consistent with this hypothesis, CT cells in lower L6 have more vM1 presynaptic partners than those in upper L6 (Whilden et al., 2021).

To explore these questions, we examined the mechanisms whereby CT cells with distinct intrathalamic axonal projections are engaged by vM1 input. For this, we combined optogenetic strategies and retrograde labeling techniques to stimulate vM1 afferent selectively and record from identified CT neurons. Our results reveal the cellular and synaptic mechanisms by which a subclass of L6 CT cells projecting to both VPm and posterior medial nucleus (POm) are engaged by vM1 and suggest a novel view of how a subcircuit of CT neurons in vS1 may influence thalamic information processing during active touch.

## MATERIALS AND METHODS

### Animals

All procedures were carried out in accordance with the National Institutes of Health (NIH) Guidelines *for the Care and Use of Laboratory Animals* and approved by the Institutional Animal Care and Use Committee (IACUC) at Michigan State University. We used the following mouse lines in this study: *Ntsr1-Cre* (catalog # 017266-UCD, Mutant Mice Research & Resource Center; RRID: MMRC_017266-UCD); (Gong et al., 2007), Ai14 (catalog # 007908, The Jackson Laboratory; (Madisen et al., 2010), and Crl:CD1 (ICR) (catalog # 022, Charles River Laboratories). Homozygous *Ntsr1-Cre* mice were crossed with homozygous Ai14 reporters or ICR mice, resulting in heterozygous experimental animals for the indicated genes. We occasionally used the following homozygous Cre driver mice crossed to ICR mice for thalamic injections: *PV-Cre* (catalog # 008069, The Jackson Laboratory: 008069; (Hippenmeyer et al., 2005) and *Grp-Cre* (catalog # 031183, Mutant Mice Regional Resource Center; RRID:MMRRC_031183-UCD) (Gerfen et al., 2013). All mouse strains used in the study had Crl:CD1 genetic backgrounds. Experimental animals were group-housed with same-sex littermates in a dedicated animal care facility maintained on a 12:12 h light/dark cycle. Food and water were available *ad libitum*. We used both male and female mice in this study.

### Stereotaxic injections

For stereotaxic injections, ∼4-week-old mice were used (Mean injection age: 27.2 ± 3.1 days, range 20 – 35 days). All surgeries were performed as previously described (Crandall et al., 2017; Martinetti et al., 2022). Briefly, mice were anesthetized with a Ketamine-Dexdomitor cocktail diluted in sterile saline (KetaVed, 70-100 mg/kg; Dexdomitor, 0.25 mg/kg; intraperitoneally), and once deeply anesthetized, placed into a stereotaxic frame with an integrated warming base that maintained constant core body temperature (Stoelting). Next, we placed a thin layer of ophthalmic ointment over the eyes to prevent damage or drying (Patterson Veterinary Artificial Tears), made a scalp incision with fine surgical scissors, and retracted the scalp and periosteum to expose the skull. A small craniotomy and durotomy were then performed, and a glass micropipette was loaded with either a virus or tracer and lowered slowly into the brain. A small volume of virus or tracer was then pressure-ejected (Picospritzer pressure system) over 5-10 min. Afterward, the pipette was held in place for 5-10 min before slowly being withdrawn from the brain. Following the injection(s), the incision was closed with a surgical adhesive (GLUture), and a local anesthetic (Bupivacaine; 2 mg/kg) and anti-sedative (Antisedan; 2.5 mg/kg) were administered to all animals. All mice recovered on a heating pad for ∼1 hr before returning to their home cage.

To optically stimulate vM1 axons/terminals, we unilaterally injected an adeno-associated virus (volume: 0.1-0.2 μL per injection site; titer = 3.1-3.5 x 10^12^ viral genomes/mL) that encoded genes for hChR2 (H134R)-EYFP fusion proteins (rAAV2/hSyn-hChR2[H134R]-eYFP-WPREpA; lot #AV4384H, University of North Carolina Viral Vector Core) into the right vM1 (Martinetti et al., 2022) (vM1 coordinates relative to bregma: 0.9 and 1.3 mm anterior, 1.25 mm lateral, and 0.4 and 1.0 mm ventral from surface). We previously defined the center of M1 anatomically to be between 0.9 mm and 1.3 mm anterior and 1.25 mm lateral with respect to bregma (Martinetti et al., 2022), which is consistent with studies defining vM1 in the mouse anatomically and with intracortical microstimulation (Ferezou et al., 2007; Mao et al., 2011; Sreenivasan et al., 2016). We chose AAV2 because the diffusion is more confined than other serotypes (Aschauer et al., 2013; Haery et al., 2019), allowing for more precise vM1 targeting and limiting lateral spread (Martinetti et al., 2022). All injections were done at two depths, corresponding to superficial and deep layers in vM1 (Hooks et al., 2011; Mao et al., 2011), and two anterior-posterior locations, in agreement with the narrow region vM1 occupies in the frontal cortex (Ferezou et al., 2007; Mao et al., 2011; Sreenivasan et al., 2016).

To retrogradely label L6 CT cells, we unilaterally injected both the right POm and VPm simultaneously with Cholera toxin-B conjugated to either Alexa 647 (CTB-647, ThermoFisher C34778) or Alexa 555 (CTB-555, ThermoFisher C34776). Both CTB solutions were prepared by dissolving in 0.1 M phosphate-buffered saline to a final injection concentration of 1.0 μg/μL (volume: 100-150 nL per nucleus). In a subset of control experiments, we injected a latex microsphere-saline mixture (red Retrobeads, Lumafluor, Cat# R180) into the VPm (volume: 100-150 nL; 1:40 dilution) (Coordinates relative to bregma, VPm: 1.25 mm posterior, 1.8 mm lateral, and 3.10 mm ventral from the surface; POm: 1.35 mm posterior, 1.4 mm lateral, and 2.87 mm ventral from the surface). The mean expression time following each injection was 20.9 ± 0.3 days, range 13 – 29).

### Acute brain slice preparation

After ∼3 weeks of viral and tracer expression, acute brain slices containing vS1 ipsilateral to the vM1 injection were prepared for *in vitro* recording and imaging, as previously described (Wundrach et al., 2020; Martinetti et al., 2022). Briefly, mice (postnatal day 37-55, mean = 46 days) were deeply anesthetized using isoflurane, decapitated, and then their brain rapidly extracted and placed in a cold (∼4 °C) oxygenated (5% CO_2_, 95% O_2_) slicing solution containing the following (in mM): 3 KCl, 1.25 NaH_2_PO_4_, 10 MgSO_4_, 0.5 CaCl_2_, 26 NaHCO3, 10 glucose, and 234 sucrose. Acute coronal brain slices (300 µm thick) were then sectioned (VT1200s, Leica) and transferred to a holding chamber filled with warm (32 °C) oxygenated artificial cerebrospinal fluid (ACSF) containing (in mM): 126 NaCl, 3 KCl, 1.25 NaH_2_PO_4_, 2 MgSO_4_, 2 CaCl_2_, 26 NaHCO_3_, and 10 glucose. Slices were incubated at 32 °C for 20 minutes and then at room temperature for at least 40 minutes before recording. We collected brain slices containing vM1, vS1, POm, and VPm for subsequent imaging to assess injection accuracy, tissue health, and expression patterns. We only considered mice with no apparent signs of tissue damage from the craniotomy or off-target injections. Our analysis of targeting the POm and VPm was three-fold. First, we distinguished VPm from POm in living slices cut in the coronal plane through the somatosensory thalamus. Under brightfield transillumination in the submersion chamber, the POm appears as a lighter triangle-shaped region medial and dorsal to the ovoid-shaped VPm that, in turn, is medial to the ventral posterior lateral nucleus (VPl) and thalamic reticular nucleus (TRN) (Jones, 2007; Landisman and Connors, 2007) (Fig. 3*A* and Fig. 3-1*A*). Second, we compared the overlap between ChR2-EYFP-expressing M1 axons, which target the POm but not the VPm (Yamawaki and Shepherd, 2015), and the two spectrally distinct retrograde tracers (Fig. 3-1*A*). Lastly, we used the anterograde labeling properties of CTB to inspect the cortical projection patterns of each injection site (Dederen et al., 1994; Abbott et al., 2013) and compared those patterns to the known termination patterns of VPm and POm cells, which target primarily L4 and L5a/L1, respectively (Herkenham, 1980; Wimmer et al., 2010; Crandall et al., 2017) (Fig. 3*B*). Occasionally, the CTB spread to the adjacent nucleus, which was easily recognizable by the criteria listed above. These mice were excluded from this study.

**Figure 3-1.**
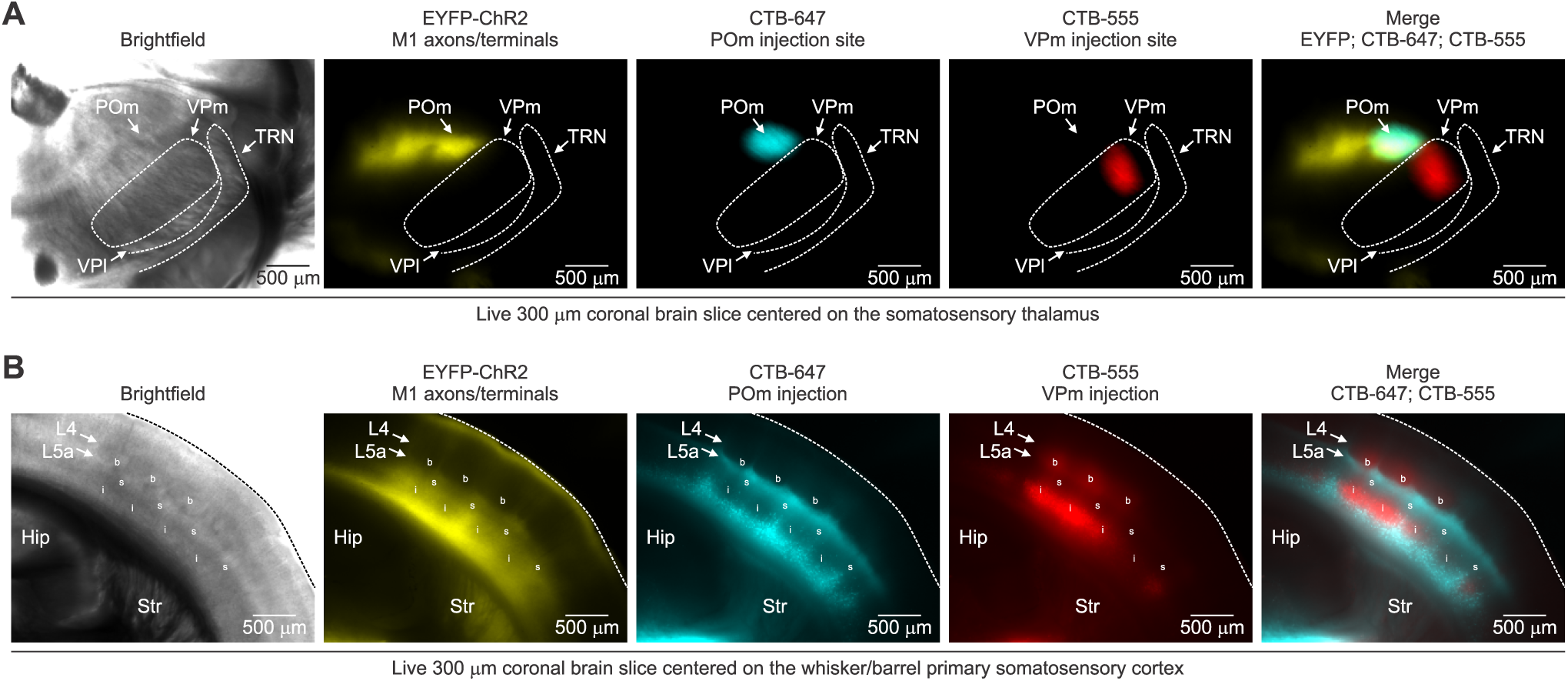
Confirming POm targeting based on M1 projections to the POm and the distribution of retrogradely labeled CT cells in L6a. ***A,*** Left, Brighfield image of a live coronal slice (300 μm) through somatosensory thalamus S1. Right, epifluorescence images of the same slice showing M1 axons/terminals in the thalamus (EYFP/ChR2) and the dual CTB injections (CTB-647 and CTB-555) into the POm and VPm. Note the overlap of EYFP and CTB-647 in the POm, confirming a successful hit. ***B,*** Left, Brighfield image of a live coronal slice (300 μm) through vibrissal/barrel primary somatosensory cortex (S1). Right, epifluorescence images of the same slice showing M1 axons/terminals in the S1 and distribution of retrogradely labeled cells from the POm (CTB-647) and VPm (CTB-555). POm, posterior medial nucleus; VPm, ventral posterior medial nucleus; VPl, ventral posterior lateral nucleus; TRN, thalamic reticular nucleus; Hip, hippocampus; Str, striatum; b, L4 barrel, s, septa; I, L6a infrabarrel.

### In vitro electrophysiological recordings and data acquisition

Brain slices were transferred to a submersion chamber on an upright microscope (Zeiss Axio Examiner.A1) and continually superfused with warm (32 ± 1 °C) oxygenated ACSF containing (in mM): 126 NaCl, 3 KCl, 1.25 NaH_2_PO_4_, 1 MgSO_4_, 1.2 CaCl_2_, 26 NaHCO_3_, and 10 glucose. Neurons were visualized using infrared differential interference contrast (IR-DIC) and fluorescence microscopy using a 40x water-immersion objective (Zeiss, W Plan-Apo 40x/1.0 NA) and a video camera (Olympus, XM10-IR). Whole-cell current-clamp recordings were obtained using patch pipettes (4-6 MΩ) pulled (Sutter P-1000, Sutter Instruments) from filamented borosilicate capillary glass (Sutter Instruments) and back-filled with a potassium-based internal solution containing (in mM): 130 K-gluconate, 4 KCl, 2 NaCl, 10 HEPES, 0.2 EGTA, 4 ATP-Mg, 0.3 GTP-Tris, and 14 phosphocreatine-K (pH 7.25, 290-300 mOsm). For all voltage-clamp experiments, pipettes contained the following cesium-based internal solution (in mM): 130 Cs-gluconate, 4 CsCl, 2 NaCl, 10 HEPES, 0.2 EGTA, 4 ATP-Mg, 0.3 GTP-Tris, and 14 phosphocreatine-K (pH 7.25, 290-300 mOsm). All voltages were corrected for a -14 mV liquid junction potential. Electrophysiological data were acquired and digitized at 20 kHz (50 kHz when assessing single action potential properties; MultliClamp 700B, Digidata 1550B4, pClamp 11), and low-pass filtered at 3 kHz (voltage-clamp) or 10 kHz (current-clamp) before digitizing. The pipette capacitances were neutralized, and the series resistances (typically 10-25 MΩ) were compensated online (70 – 80% for voltage-clamp and 100% for current-clamp). Series resistances were continually monitored and adjusted online to ensure accurate compensation.

### Photostimulation

ChR2-expressing axons/terminals were stimulated optically using a high-power white light-emitting diode (LED; Mightex, LCS-5500-03-22) driven by an LED controller (Mightex, BLS-1000-2). LED on/off times were fast (<50 µs) and had a constant amplitude and duration, verified by a fast photodiode (Thorlabs, DET36A). Collimated light (0.5 ms flashes) was reflected through a single-edge dichroic beam-splitter (Semrock, FF660-FDi02) and a high-magnification water-immersion objective (Zeiss, W Plan-Apo 40x/1.0 NA), with an estimated spot diameter of ∼1500 µm, and a maximum LED power of ∼20 mW (∼11.3 mW/mm^2^) used in experiments. When comparing subthreshold synaptic responses, the LED power was typically ∼8 mW (∼4.5 mW/mm^2^), or 4x threshold. For all recordings, photostimulation was performed over the distal segments of vM1 axons in vS1 and centered over the somata of the recorded neurons. When recording from Ntsr1;Ai14 mice (Fig. 2), we adjusted the LED intensity to 1-6x the threshold for evoking a synaptic response in each neuron when held at their preferred resting membrane potential (RMP; ∼ -84 mV) in current-clamp. For all paired CT recordings (Figs. 4 and 6), we adjusted the LED intensity to 1-6x the threshold for evoking a synaptic response in the Dual CT neuron when held at their RMP in current clamp (potassium internal solution) or near the reversal potential for inhibition (-79 mV) and excitation (∼0 mV) in voltage clamp (cesium internal solution).

### Electrophysiological data analysis

We performed electrophysiological data analysis using Molecular Devices Clampfit 11 and Microsoft Excel. Synaptic responses to optical stimulation were measured from postsynaptic neurons in whole-cell current- and voltage-clamp. The evoked EPSC/EPSP/IPSC amplitude was measured relative to a baseline before the stimulus ( 5 ms). The EPSC/EPSP and IPSC peak search windows were set to 10 ms and 20 ms immediately after the onset of the stimulus, respectively. Values were based on mean responses to 2-10 stimuli (typically 5). Synaptic latencies were measured from stimulus onset. We used previously described methods to analyze the intrinsic membrane properties of neurons (Crandall et al., 2017). L6 CT cells were identified and targeted for recording in two ways: 1) by their expression of the red fluorescent protein (tdTom) in Ntsr1-Cre;Ai14 mice or 2) by their retrograde labeling from POm, VPm, or both. All VPm-only projecting CT neurons targeted for recording were exclusively located in the upper portion of L6a (L6a_U_; 70 – 84% depth from pia), and all Dual VPm/POm projecting CT neurons targeted for recording were exclusively located in the lower portion of L6a (L6a_L_; 84 – 97% depth from pia). These laminar borders are consistent with previous reports for L6 in vS1 (Frandolig et al., 2019; Martinetti et al., 2022). We excluded recordings from cells whose soma were not located at the specified depths. We also excluded cells in L6b from our analysis (L6b: 97 – 100% depth from pia).

### Immunohistochemistry and tissue processing for cell counts

All tissues were prepared from acute coronal brain slices, as described previously (Martinetti et al., 2022). Briefly, acute brain slices (300 µm thick) containing vS1 were fixed by placing in a 4% paraformaldehyde in 0.1 M phosphate-buffered saline (PBS) solution overnight at 4°C. Slices were then transferred to a 30% sucrose in 0.1 M PBS solution and stored at 4°C until re-sectioning (2-3 days). The tissue was then re-sectioned at 50 µm using a freezing microtome (Leica, SM2010R) and immunostained for the neuronal nuclear antigen, NeuN (Mullen et al., 1992), using methods described previously (Crandall et al., 2017). Sections were washed twice in 0.1 M PBS followed by 3 washes in 0.1 M PBS with 0.15 M NaCl (5 min/wash). After washing, the sections were incubated at room temperature for 1 hr in a blocking solution containing 0.1% Tween, 0.25% Triton X-100, 10% normal goat serum in PBS. Sections were then incubated with primary antibody for 3-4 days in a rotating platform at 4°C. Following primary incubation, the tissue was washed 5 times in PBS (5 min/wash), then pre-blocked in a blocking solution (same as above; 45 min), incubated with a secondary antibody for 2 hrs at room temperature, and washed 3 times in PBS (10 min/wash). All sections were placed on a glass slide and cover-slipped using Vectashield with DAPI (Vector laboratories H-1200). The primary antibody used was a mouse monoclonal anti-NeuN (1:1000; Millipore MAB377), and the secondary antibody used was a goat anti-mouse IgG (H+L) with either an Alexa Fluor 488 (1:250; Invitrogen A11029) or Alexa Fluor 647 congugate (1:250; Invitrogen A21236).

### Imaging and cell counts

Sections processed for immunohistochemistry were initially visualized using an epifluorescent microscope (Zeiss Axio Imager.D2) and Zen software to identify sections for cell counting. Selection criteria for a single barrel column included the following: at least 3 adjacent L4 barrels were anterogradely filled by VPm terminals (due to CTB anterograde properties) (Dederen et al., 1994; Abbott et al., 2013), each barrel exhibited L5a “cupping” by anterogradely filled POm terminals, and the aligned L6 expressed retrogradely filled CT cells. Selected sections were then visualized on a confocal microscope (Nikon A1 Laser Scanning Confocal Microscope) using a 40x (1.0 N.A.) or 100x (1.49 N.A.) and laser excitation 405nm, 488nm, 561nm, and 647nm. Z-stacks of the full depth of the section were obtained at 0.8 µm/plane.

For each section, we imaged across the vertical extent of the cortex, from the middle of L4 to the bottom of white matter. The transition between L4 and L5a was used as the upper boundary, and the white matter as the lower boundary when normalizing somatic position across preparations. The upper and lower borders of L6a were determined by measuring the CTB fluorescence representing retrogradely-filled VPm CT neurons. We focused our L6 CT cell counts on a narrow 120 μm region across L6 that corresponded to the middle of the L4 barrel and did not include septa. Colocalization of both NeuN and retrograde tracers was quantified using the Cell Counter plugin in Fiji (Schindelin et al., 2012).

### Experimental Design and Statistical Analysis

In this study, we used no formal method for randomization, and the experimenters were not blind to experimental groups during data collection or analysis. No statistical methods were used to identify the sample size, but our sample size is similar to those previously reported (Crandall et al., 2017; Martinetti et al., 2022). Statistical analyses were performed using OriginPro software. We used a Shapiro-Wilk test to determine whether the data had been drawn from a normally distributed population. Parametric tests were used for normally distributed populations, whereas non-parametric tests were used if a normality assumption was invalid. Significance was assessed using the appropriate parametric (Paired t-test or two-sample t-test) or non-parametric test (Wilcoxon paired sign-rank test or Mann-Whitney U test), as indicated in the results. All tests were two-tailed. Statistical significance was defined as p < 0.05, and data are represented as mean ± SEM.

## RESULTS

### vM1 preferentially targets corticothalamic circuits in lower L6a of vS1

To investigate how vM1 projections engage L6 CT circuits in vS1, we combined *in vitro* electrophysiological recordings with optogenetic control strategies (Crandall et al., 2017; Martinetti et al., 2022). To achieve selective optical activation of the vM1 pathway, we expressed the light-gated cation channel Channelrhodopsin-2 fused to an enhanced yellow fluorescent protein (ChR2-EYFP) in vM1 neurons and subsequently their axons/terminals in vS1 by viral transduction (Fig. 1*A*). Viral injections were performed unilaterally in vM1 of *Ntsr1-Cre* mice, which expresses Cre-recombinase in nearly all L6 CT neurons of sensory cortex (Gong et al., 2007; Bortone et al., 2014; Kim et al., 2014; Guo et al., 2017), crossed with Cre-dependent tdTomato (tdTom) reporters (Ai14) (Madisen et al., 2010). Three weeks after injections, we found intense ChR2-EYFP-expressing vM1 axons concentrated in L1 and L5/6 of vS1, consistent with previous reports (Veinante and Deschenes, 2003; Rocco and Brumberg, 2007; Lee et al., 2013; Kinnischtzke et al., 2014; Martinetti et al., 2022). However, we also observed dense, uniform labeling of vM1 axons/terminals in the lower portion of L6a (L6a_L_) within vS1 and weak, discontinuous labeling in the upper part of L6a (L6a_U_) (Fig. 1*A*, *B*). The spatial organization of vM1 projections in L6a_U_ had a columnar appearance resembling infrabarrels (Crandall et al., 2017), with clear labeling in the septa and considerably less located within the center of L6 infrabarrels (Fig. 1*B*). Averaging the fluorescence intensity profiles from multiple slices and measuring the mean fluorescence across L6a sublamina revealed vM1 axons/terminals were denser in L6a_L_ than L6a_U_ (Figs. *1C*). We observed minimal fluorescence in L6b, a distinct layer of neurons with different gene expression patterns, morphology, connectivity, and perhaps functions (Heuer et al., 2003; Chen et al., 2009; Marx and Feldmeyer, 2013; Hoerder-Suabedissen and Molnar, 2015; Hoerder-Suabedissen et al., 2018; Zolnik et al., 2020; Zolnik et al., 2024). These fluorescence profiles suggest that vM1 inputs could make strong synaptic connections onto CT cells in L6a_L_ of vS1.

**Figure 1.**
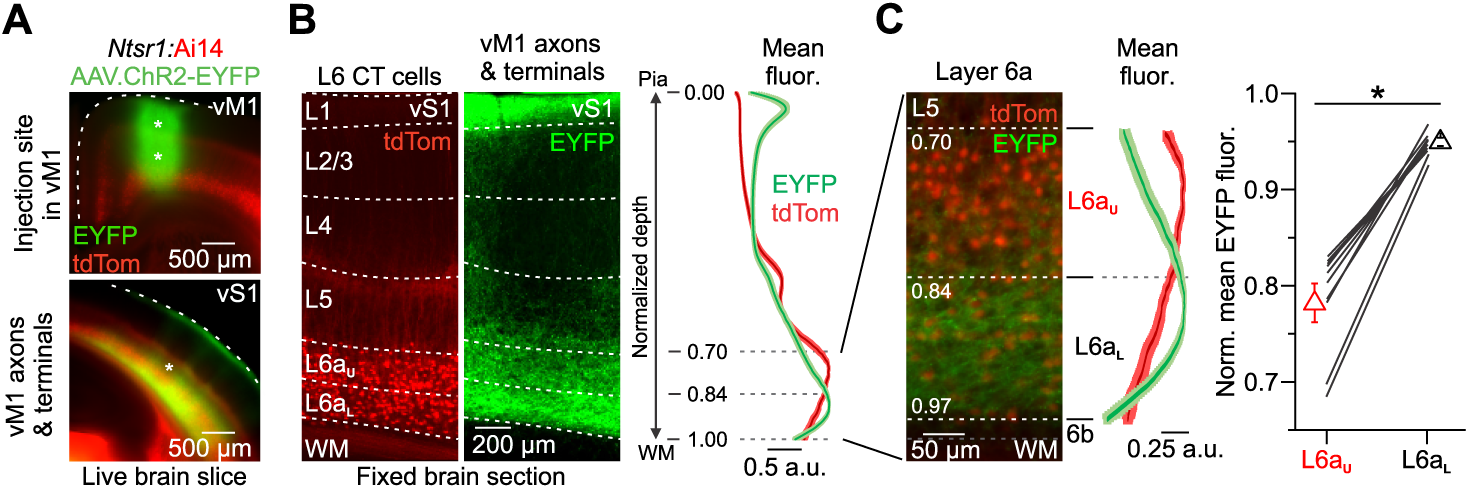
vM1 axons/terminals define two L6a sublayers in vS1. ***A,*** Top, Fluorescent image of a live coronal brain slice (300 µm) taken from an *Ntsr1-Cre* x floxed-tdTom reporter mouse (Ai14) expressing ChR2-EYFP in vM1 at the viral injection sites (asterisks). Bottom, Fluorescent image showing ChR2-EYFP-expressing vM1 axons/terminals in vS1 of the same mouse. ***B,*** Left and middle, High-magnification fluorescent image of a 50-µm-thick section obtained from the same vS1 slice in ***A*** (asterisk in the lower image), showing tdTom-expressing L6 CT cells and EYFP-labeled vM1 axons/terminals terminating in vS1. Right, Comparison of the normalized CT cell (red) and vM1 axons/terminal (green) fluorescence as a function of depth from the pial surface. ***C,*** Left, High-magnification fluorescent image of the same slice shown in ***B*** with corresponding fluorescence as a function of depth through L6. Upper and lower L6a were defined based on the normalized depth from pia (L6a_U_ from 70 to 83%; L6a_L_ from 84 to 97%). Right, Summary plot of the normalized mean EYFP fluorescence in L6a_U_ and L6a_L_ of vS1 in the same slice (Mean normalized fluorescence: L6a_U_: 0.78 ± 0.02 a.u., L6a_L_: 0.95 ± 0.00 a.u.; *n* = 10 slices, 10 mice, *p* = 0.00592, Wilcoxon signed-rank test). The asterisk indicates statistical significance. Data are represented as mean ± SEM.

Since anatomical colocation of axon/terminal and soma/dendrites is not always a great predictor of synaptic strength, we next sought to understand how vM1 afferents engage CT circuits by performing targeted current-clamp recordings from tdTom-expressing cells throughout the depth of L6a in vS1. We initially tested the impact of vM1-evoked synaptic responses while recording CT cells at their resting membrane potential in current-clamp (RMP: ∼-84 mV) (Fig. 2*A*). Brief (0.5 ms) widefield optical stimulation of vM1 inputs evoked short latency (<2 ms) excitatory postsynaptic potentials (EPSPs) that never drove action potential (AP) firing in any L6a CT cell recorded from RMP across a wide range of stimulus intensities (1-6x EPSP threshold; L6a_U_: 0 of 19 cells, 4 mice; L6a_L_: 0 of 21 cells, 4 mice). However, despite not driving L6 CT cells to spike from RMP, we did observe that subthreshold EPSPs were, on average, larger in L6a_L_ CT neurons than L6a_U_ CT cells (6x intensity EPSP peak: L6a_L_: 9.1 ± 1.0 mV; L6a_U_: 4.2 ± 0.6 mV; *p* = 1.78e-4, two-sample t-test; Fig. 2*A*, right).

**Figure 2.**
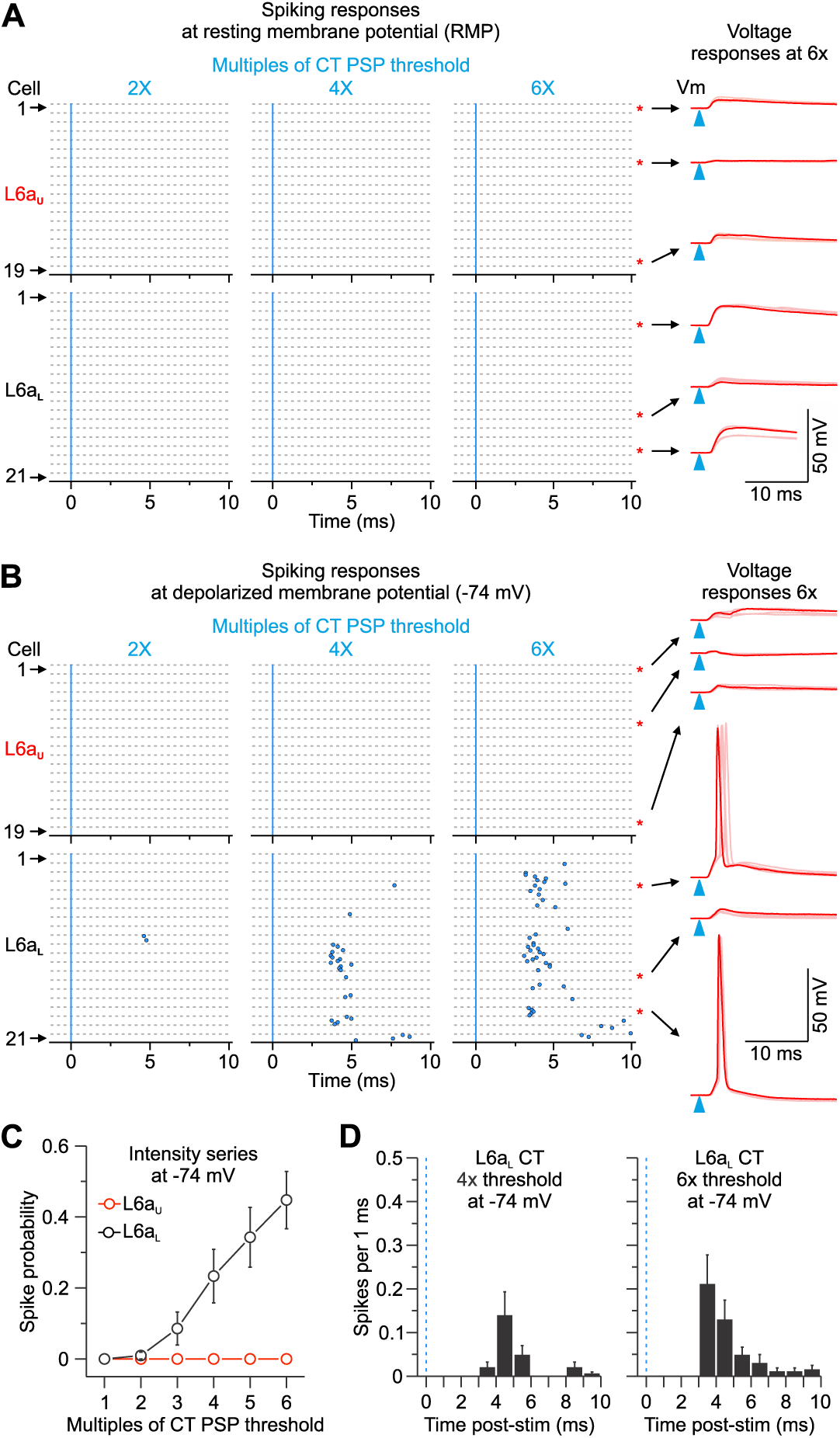
vM1 inputs differentially activate CT populations in L6a of vS1. ***A,*** Left, Raster plots of L6 CT cell spiking (blue circles), sorted by L6a_U_ and L6a_L_, evoked by single LED flashes (blue line; 0.5 ms) of increasing LED intensities while holding each cell at resting membrane potential (RMP). For each cell, individual trials (5 trials/cell) are plotted as clustered groups between grey lines for display. Stimulus intensities normalized to the EPSP threshold for each cell. The black dashed line indicates the border between L6a_U_ and L6a_L_. Right, vM1 synaptic responses in response to a 6x threshold light intensity from 6 different CT cells at RMP (overlay of 5 sweeps shown). The blue triangles show LED stimulation. ***B,*** Same as ***A*** above, but each cell is held at a depolarized membrane potential (-74 mV) with injected current. ***C,*** Summary of population spike probability at -74 mV with increasing LED stimulus intensities for L6a_U_ (*n* = 19 cells, 5 mice) and L6a_L_ CT cells (*n* = 21 cells, 5 mice) (*p* = 6.48e-12, two-way ANOVA). ***D,*** Population peristimulus time histograms (PSTHs) plotting spike probability per 1 ms bin at -74 mV (Light intensities = 4x and 6x EPSP threshold) for all L6a_L_ CT cells recorded (*n* = 21 cells from 5 mice). The blue line indicates the onset of the LED. Values are represented as mean ± SEM.

Although cells did not spike from RMP, studies have shown L6 CT neurons to be active in awake animals engaged in behaviors (Swadlow and Weyand, 1987; Briggs and Usrey, 2009; Augustinaite and Kuhn, 2020; Voigts et al., 2020; Clayton et al., 2021; Dash et al., 2022), including those in vS1 during whisking and locomotion (Dash et al., 2022), which are behaviors associated with depolarization of the membrane potential for many cortical neurons (Steriade et al., 2001; Crochet and Petersen, 2006; McGinley et al., 2015). To characterize the functional impact of vM1 input under this more active state, we injected individual CT cells with positive current to produce a steady depolarization of -74 mV at the soma (∼10 mV more depolarized than RMP) (Fig. 2*B, C*). In contrast to responses at RMP, vM1 input evoked AP firing in most L6a_L_ CT cells (16 of 21 cells at 6x; 76%;) with short spike latencies consistent with monosynaptic drive (4x intensity: 5.4 ± 0.5 ms, *n* = 8 cells; 6x intensity: 5.3 ± 0.5 ms, *n* = 16 cells); Fig. 2*D*). However, membrane depolarizations were insufficient to facilitate spiking in any L6a_U_ CT cells (0 of 19 cells at 6x; 0%;) over a range of intensities (Fig. 2*C*). These findings indicate that vM1 inputs differentially engage CT cells within vS1 based on their anatomical position within L6.

### Consistent with their spatial distribution, Dual CT cells respond strongly to vM1 input

Our data suggest that among the CT neurons in L6a of vS1, vM1 provides the strongest excitatory inputs to cells in the lower half of the layer, L6a_L_. Thus, we asked whether the strength of vM1 input correlates with the specific class of CT neuron. Indeed, in rodent vS1, L6 CT cells can be classified into two broad subclasses based on their intrathalamic axonal projections (Bourassa et al., 1995; Killackey and Sherman, 2003; Chevee et al., 2018; Whilden et al., 2021). The first class is located in L6a_U_ and projects primarily to the VPm, whereas the other is situated in L6a_L_ and projects to the POm, with many also collateralizing in VPm. However, whether all POm-projecting CT cells target VPm and the relative contribution of each class to the total CT population in the upper and lower half of L6a is unclear.

To directly test these questions and, more importantly, to label distinct subclasses of L6a CT neurons for subsequent recordings, we stereotactically injected inert and spectrally distinct retrograde tracers (CTB-647 and CTB-555) in the VPm and POm of the same mouse and analyzed the distribution of labeled cells (Fig. 3*A*). After allowing ∼3 weeks for retrograde transport to the cell body, we observed fluorescent labeling of a large population of CT cells in L6, revealing a visible sublamina distribution, with the distribution of labeling from the VPm mostly observed in L6a_U_ and labeling from the POm primarily found in L6a_L_ (Fig. 3*B*). Consistent with previous observations, we occasionally observed more significant labeling from the VPm within L6a_U_ infrabarrels positioned directly under L4 barrels (Crandall et al., 2017), and more labeling from the POm within the L6a_U_ septa (Killackey and Sherman, 2003) (Fig. 3-1*B*). Interestingly, closer inspection of individual retrogradely labeled cells revealed 3 distinct subclasses of CT neurons based on the presence of 1 or 2 retrograde tracers in their soma and thus their axonal projection patterns: VPm-only projecting, POm-only projecting, and Dual projecting, whose axons target both VPm and POm (Fig. 3*C*). Overall, we found that labeled CT cells represented 58.3 ± 0.9% of the total neuronal population in L6a of mouse vS1 (7 columns, 4 mice; Table 1), consistent with previous reports (Bortone et al., 2014; Kim et al., 2014; Crandall et al., 2017). We also found that 93.0 ± 0.9% of all labeled CT neurons in L6a projected to the VPm, with most identified as VPm-only projecting (51.0 ± 4.6%) and the remaining classified as Dual projecting (42.0 ± 5.4%). In contrast, we found very few POm-only projecting CT cells in L6a (7.0 ± 0.9%). These results validate previous assumptions that most CT cells that project to the POm also collateralize in the VPm (83.7 ± 4.2%). Furthermore, comparing the distribution of labeled cells revealed L6a_U_ is populated mainly by VPm-only CT cells (78.1 ± 6.4%), while L6a_L_ is populated primarily by Dual projecting CT neurons (72.2 ± 6.8%; Fig. 3*D*). Although we found all CT subclasses in L6b, they represented a smaller percentage of the total population (29%; 12 of 42 cells, 7 columns, 5 mice). Thus, these data indicate that most CT cells in L6a of vS1 can be grouped into two broad subclasses based on their intrathalamic projection targets and sublamina position, with L6a_U_ populated primarily by VPm-only projecting CT cells and L6a_L_ populated mainly by Dual projecting CT neurons.

**Figure 3:**
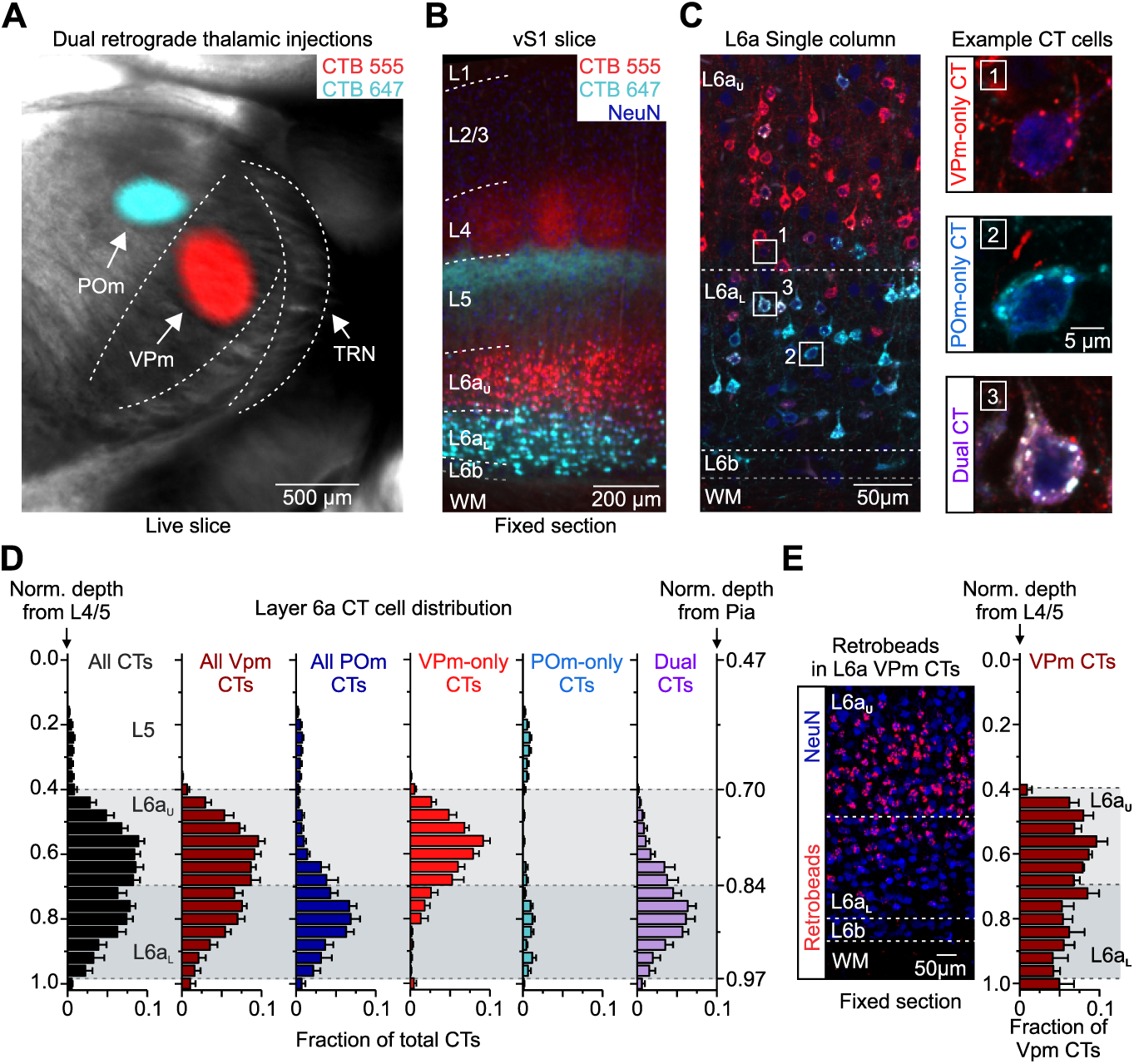
Distribution within L6a of 3 classes of CT cells based on their axonal projection patterns. ***A,*** Live slice (300 µm) image through the somatosensory thalamus taken from a mouse injected with spectrally distinct retrograde tracers in the VPm (CTB-555, red) and POm (CTB-647, cyan). Shown is an overlay of fluorescence with brightfield. ***B,*** Low-magnification fluorescent image of a 50-µm-thick section through vS1 taken from the live slice shown in ***A***. Tissue was stained immunohistochemically for NeurN (blue). ***C,*** Left, Confocal image of L6 taken from the same section shown in ***B***. Right, High-magnification images of the boxed areas in the left image. Shown are three types of CT neurons identified by their expression of one or two fluorescent retrograde tracers: VPm-only (top: project only to the VPm), POm-only (middle: project only to the POm), and Dual CT cells (bottom: project to both VPm and POm). ***D,*** Normalized distribution histograms of CTB-labeled CT cells based on their intrathalamic axonal projections (*n* = 1,294 cells in L6a and L6b, 7 barrel columns, 5 mice). Cell depths were normalized to the L4/5 border (left axis), but the normalized depth from pia is also shown (right axis). L6a_U_ was defined as 0.42 ± 0.01 to 0.69 ± 0.01 (light gray area), while L6a_L_ was defined as 0.69 ± 0.01 to 0.97 ± 0.02 (darker gray area). Many VPM-only CT neurons are found in L6a_U_, whereas many Dual CT cells are located in L6a_L_. ***E,*** Left, Confocal image of a 50-µm-thick section through L6 of vS1 taken from a mouse injected with red Retrobeads in VPm. Tissue was stained immunohistochemically for NeurN (blue). Right, Normalized distribution histogram of L6 VPm CT cells (*n* = 490 cells, 3 barrel columns, and 2 mice). CTB and retrobreads produce a similar distribution of all VPm projecting CT cells. Data are represented as mean ± SEM. See also Figure 3-1.

**Table 1.** Summary of CT neuronal population in L6a.

| Fraction of cells | All L6a | L6a<sub>U</sub> | L6a<sub>L</sub> |
| --- | --- | --- | --- |
|  | Mean ± s.e.m. | Mean ± s.e.m. | Mean ± s.e.m. |
| CT | 0.583 ± 0.009 | 0.544 ± 0.008 | 0.642 ± 0.024 |
| VPm CT | 0.930 ± 0.009 | 0.981 ± 0.007 | 0.867 ± 0.020 |
| VPm-only CT | 0.510 ± 0.046 | 0.781 ± 0.064 | 0.145 ± 0.050 |
| POm CTs | 0.490 ± 0.046 | 0.219 ± 0.064 | 0.855 ± 0.050 |
| POm-only CT | 0.070 ± 0.009 | 0.019 ± 0.007 | 0.133 ± 0.020 |
| Dual CTs | 0.420 ± 0.054 | 0.200 ± 0.061 | 0.722 ± 0.068 |

7 columns from 4 mice
\* See methods for an explanation of how cells were counted and borders measured.
\* Data shown as mean ± s.e.m.

To rule out the possibility that CTB resulted in abnormal retrograde labeling due to the possibility of it being taken up by fibers of passaged (Chen and Aston-Jones, 1995; Vercelli et al., 2000), we compared the distribution of CT cells labeled by injecting the VPm with latex microspheres (Retrobeads), which are thought not to be taken up by fibers of passage (Vercelli et al., 2000; Saleeba et al., 2019). The two modes of retrograde labeling produced a similar distribution of VPm-projecting CT cells throughout L6a, with many located in L6a_L_, where most of the POm-projecting CT cells reside (Fig. 3*E*). Therefore, the pattern of Dual CT cells labeled with CTB appears to reflect the normal distribution and proportions of this population.

To determine how vM1 engages these different subclasses of L6a CT neurons, we injected AAV-ChR2-EYFP into vM1 and spectrally distinct retrograde tracers in the ipsilateral VPm and POm of the same mouse (Fig. 4*A*, left). Subsequently, we prepared acute brain slices of vS1 to record vM1-evoked responses simultaneously or sequentially from pairs of L6a CT cells (Fig. 4*A*, right), which helps control for any variability in the level of ChR2 expression across individual slices and mice (Martinetti et al., 2022). Each pair comprised one L6a_u_ VPm-only CT cell and one L6a_L_ Dual CT neuron within the same column. In the current study, we did not record synaptic responses from POm-only CT cells because we could not rule out that these relatively few cells were mislabeled Dual CT neurons. When recording both cells in current-clamp at RMP, optogenetic stimulation of vM1 inputs evoked EPSPs that were almost always larger in Dual CT neurons than VPm-only CT cells (12 of 14 pairs, 11 mice; Fig. 4*B*), resulting in an average EPSP that was ∼3x larger in Dual CT neurons than VPm-only CT cells. This difference in vM1 responses was consistent across a wide range of LED intensities (Fig. 4*C*), suggesting that vM1 inputs preferentially target L6a_L_ Dual CT cells in vS1.

**Figure 4.**
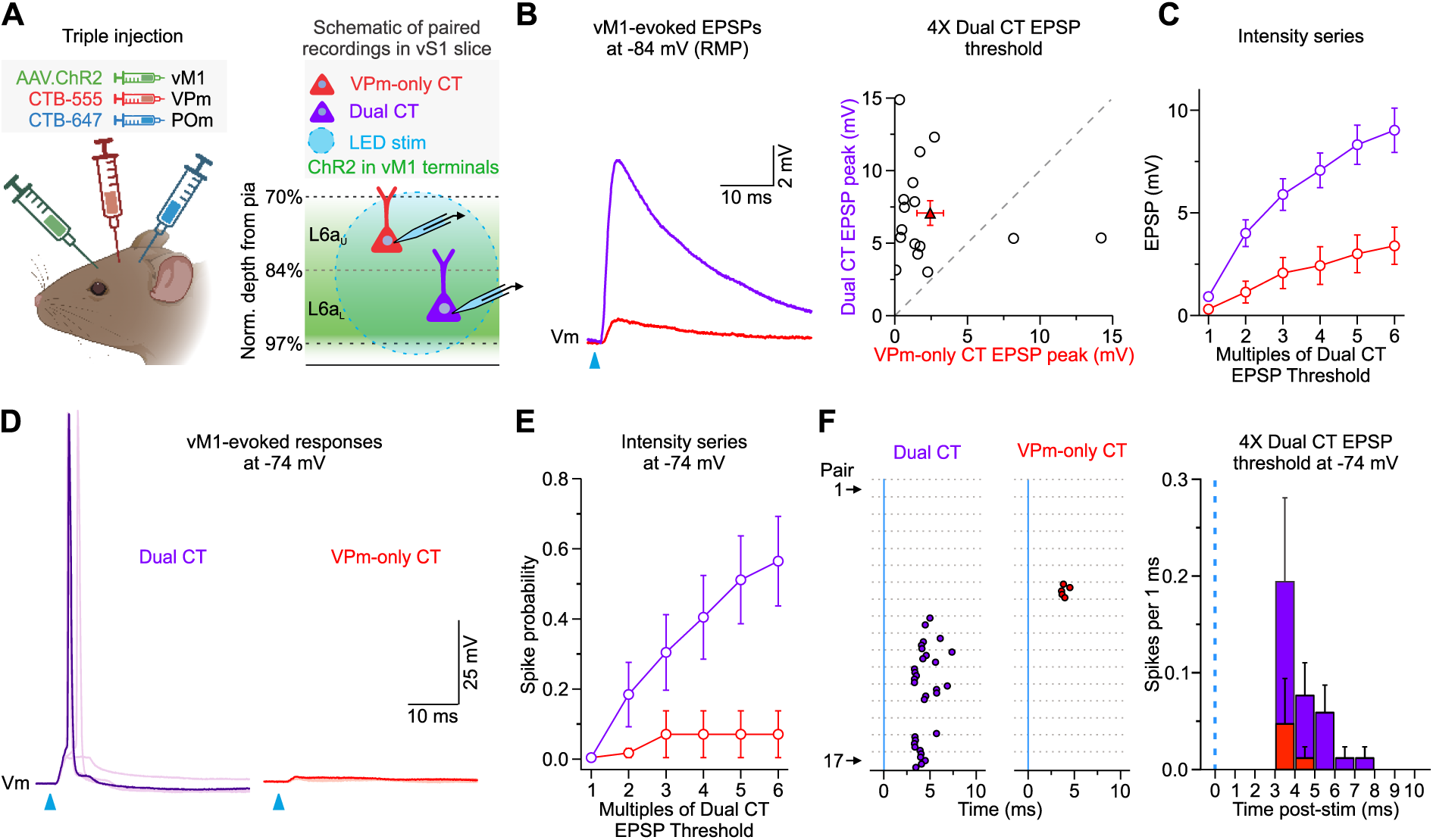
vM1 input preferentially targets L6a_L_ Dual CT neurons. ***A,*** Left, Injection schematic showing unilateral injection of AAV2-ChR2-EYFP in vM1 and spectrally distinct retrograde tracers (CTB-555 and CTB-647) in VPm and POm of the same mouse. Image created with BioRender.com. Right, Schematic of *in vitro* recording configuration showing full-field LED stimulation of ChR2-expressing vM1 axons/terminals and targeted recordings from retrogradely labeled L6a_U_ VPm-only and L6a_L_ Dual CT cell pairs. ***B,*** Left, Average vM1-evoked EPSPs from paired Dual and VPm-only CT cells when held at RMP (∼-84mV) in current-clamp. Right, Summary of EPSP peaks in response to 4x LED stimulation (Dual CT: 7.07 ± 0.85 mV; VPm-only CT: 2.43 ± 0.92 mV; *n* = 16 pairs from 11 mice; *p* = 0.00774, Wilcoxon paired sign-ranked test). ***C,*** Summary of the mean vM1-evoked EPSPs over various light intensities normalized to the Dual CT cell EPSP thresholds. Dual CTs always responded stronger than VPm-only CT neurons (*p* = 6.69e-15, two-way ANOVA, *n* = 16 pairs, 11 mice). ***D***, Example vM1-evoked responses from paired Dual and VPm-only CT cells when held at a depolarized membrane potential (-74mV) in current-clamp (overlay of 5 sweeps shown). ***E,*** Summary of population spike probability at -74 mV with increasing LED stimulus intensities (*p* = 3.54e-8, two-way ANOVA, *n* = 17 pairs, 10 mice). ***F,*** Left, Raster plots of Dual and VPm-only CT cell spiking at -74 mV evoked by single LED flashes (blue line; light intensity = 4x Dual EPSP threshold; *n* = 17 pairs,10 mice). For each cell, individual trials (5 trials/cell) are plotted as clustered groups between grey lines for display. Right, Population peristimulus time histogram (PSTH) plotting spike probability per 1 ms bin at -74 mV (Light intensities = 4x Dual EPSP threshold) for all Dual and VPm-only CT cell pairs (*n* = 17 pairs, 10 mice). The blue line indicates the onset of the LED. Values are represented as mean ± SEM.

To explore how differences in targeting impact CT cell responsiveness in a more active state, we injected each cell with positive current to produce a steady-state potential of -74 mV. Consistent with our previous recordings in *Ntsr1*;Ai14 mice (Fig. 2), at this depolarized potential, vM1 input drove AP firing in a large fraction of L6a_L_ Dual CT neurons (10 of 17 cells at 6x; 58.8%) and very few L6a_u_ VPm-only CT cells over a range of LED intensities (1 of 17 cells at 6x; 5.9%; Fig. 4*D, E*). Furthermore, the mean latency to spike for L6a_L_ Dual CT cells was consistent with monosynaptic excitatory drive (4x intensity: 4.3 ± 0.3 ms, *n* = 8 cells; Fig. 4*F*). These data indicate a sublamination of CT information processing in L6a, with Dual CT cells concentrated in the lower half, responding preferentially to vM1 input.

### Intrinsic and synaptic properties explain Dual CT responsiveness to vM1 input

Theoretically, the greater responsiveness of L6a_L_ Dual CT neurons to vM1 input could be due to their distinct intrinsic membrane properties, synaptic mechanisms, or both. Concerning intrinsic membrane properties, whole-cell current-clamp recordings revealed several differences between L6a_L_ Dual CT and L6a_U_ VPm-only CT neurons (Fig. 5*A*, Table 2). Specifically related to excitability, however, recordings showed both neurons had similar RMPs (∼-84 mV), but Dual CT neurons required ∼40% less current (rheobase) to reach AP threshold than VPm-only CT cells (Fig. 5*A-C*). Despite the lower rheobase, the voltage threshold for AP initiation was statistically similar between cell types (∼-56 mV), resulting in a similar voltage difference between rest and threshold (Fig. 5*C*). Another property that could increase the excitability of a neuron is a higher membrane input resistance. Indeed, we found that the input resistances of Dual CT neurons were ∼20% higher than those of VPm-only CT cells (Fig. 5*D, E*). POm-only CT cells had properties similar to Dual CT neurons (Table 2). These data indicate that L6a_L_ Dual and L6a_U_ VPm-only CT neurons are distinct electrophysiologically, with Dual CT cells having intrinsic membrane properties that make them more responsive than VPm-only CT neurons to afferent input.

**Figure 5.**
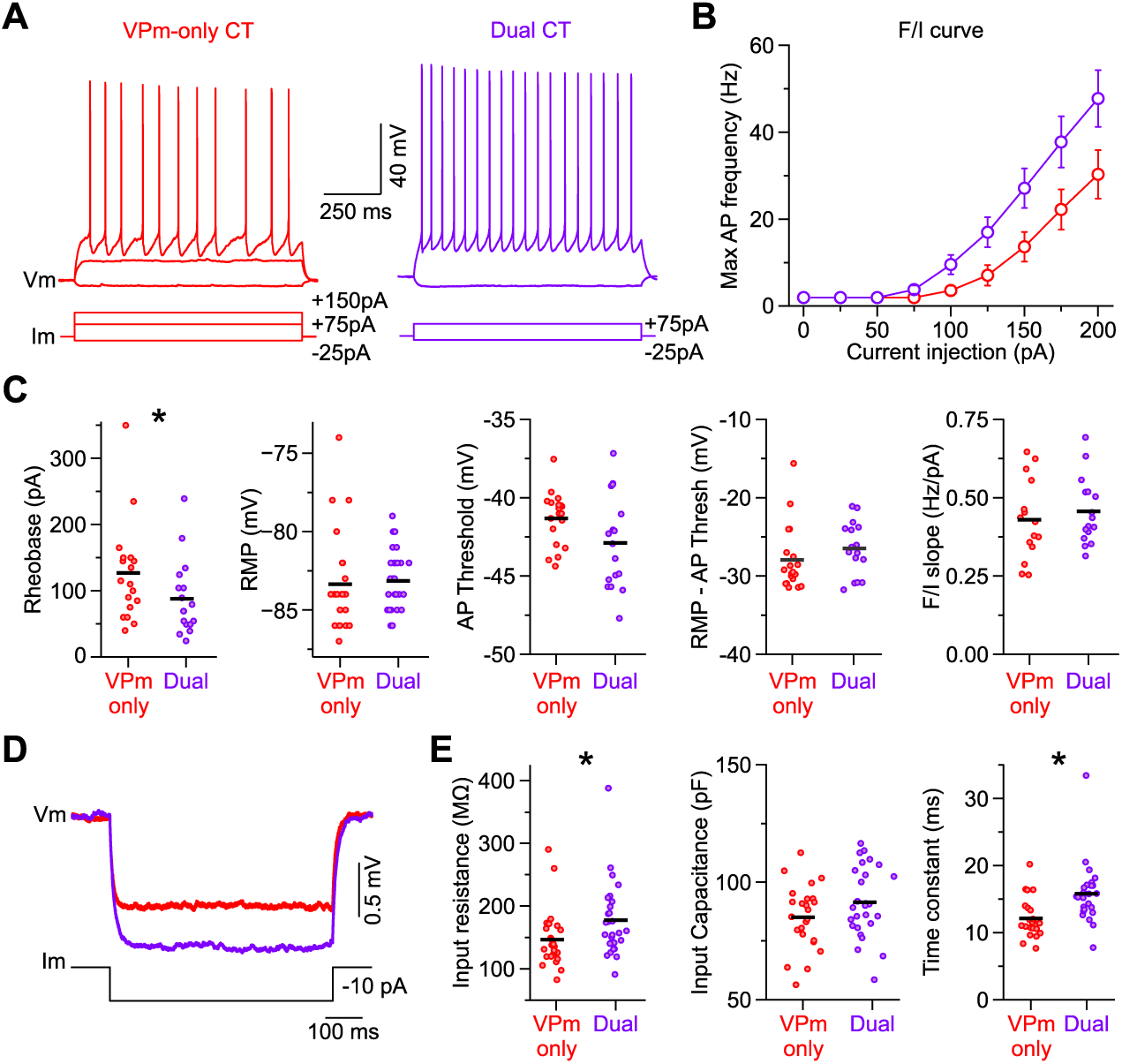
L6a CT neurons are different physiologically, with Dual CT cells being more intrinsically excitable. ***A,*** Voltage responses of VPm-only and Dual CT neurons to negative and positive current steps from RMP. The response to a positive 75 pA step was subthreshold in the VPm-only CT cell but suprathreshold for the Dual CT neuron. ***B***, Frequency-current (F/I) plot showing maximum AP frequency as a function of current injection amplitude. Data are represented as mean ± SEM. ***C,*** From left to right, summary plots of rheobase, RMP, AP threshold, the difference between RMP and AP threshold, and max F/I slope for VPm-only and Dual CT neurons. The black horizontal lines indicate the mean. ***D***, Left, Average voltage responses of VPm-only and Dual CT cells to negative current injections (-10 pA) from RMP. Responses were larger in Dual CT cells, reflecting a higher membrane resistance. ***E,*** From left to right, summary plots of membrane input resistance, input capacitance, and time constant for VPm-only and Dual CT neurons. The black horizontal lines indicate the mean. Asterisks indicate statistical significance. See Table 2 for summary statistics for all intrinsic membrane properties measured.

**Table 2.** Electrophysiological properties of retrogradely labeled CT neuronsin L6a of v51. Related to Figure 5.

|  | L6a<sub>U</sub> VPm-only CT |  |  | L6a<sub>L</sub> Dual CT |  |  | *p* | Test | L6a<sub>L</sub> POM-only CT |  |
| --- | --- | --- | --- | --- | --- | --- | --- | --- | --- | --- |
|  | Mean ± s.e.m. | *n* cells / mice |  | Mean ± s.e.m. | *n* cells / mice |  |  |  | Mean ± s.e.m. | *n* cells / mice |
| Resting membrane potential (mV) | 83.4 ± 0.6 | 25 / 13 |  | -83.1 ± 0.4 | 27 / 15 |  | 0.30538 | MW U-test | -83.6 ± 0.4 | 24 / 17 |
| Membrane resistance (MΩ) | 146.6 ± 9.2 | 25 / 13 |  | 177.5 ± 11.4 | 27 / 15 |  | 0.01210 | MW U-test | 171.8 ± 11.3 | 24 / 17 |
| Membrane time constant (ms) | 12.1 ± 0.6 | 25 / 13 |  | 15.8 ± 0.8 | 27 / 15 |  | 1.03E-04 | MW U-test | 15.7 ± 0.7 | 24 / 17 |
| Membrane capacitance (pF) | 85.0 ± 2.7 | 25 / 13 |  | 91.5 ± 2.9 | 27 / 15 |  | 0.11126 | t-test | 95.6 ± 3.8 | 24 / 17 |
| Rheobase (pA) | 126.8 ± 16.6 | 19 / 11 |  | 83.8 ± 13.6 | 17 / 10 |  | 0.02976 | MW U-test | 75.3 ± 10.3 | 18 / 14 |
| AP threshold (mV) | -55.3 ± 0.4 | 19 / 11 |  | -56.9 ± 0.7 | 17 / 10 |  | 0.05888 | t-test | -55.9 ± 0.7 | 18 / 14 |
| Rest-AP threshold (mV) | -27.9 ± 1.0 | 19 / 11 |  | -26.5 ± 0.8 | 17 / 10 |  | 0.09305 | MW U-test | -27.7 ± 0.8 | 18 / 14 |
| Delay to AP (ms) | 335.2 ± 64.0 | 19 / 11 |  | 226.8 ± 44.0 | 17 / 10 |  | 0.56842 | MW U-test | 158.3 ± 26.2 | 18 / 14 |
| AP amplitude (mV) | 85.0 ± 1.0 | 19 / 11 |  | 92.1 ± 1.7 | 17 / 10 |  | 6.17E-04 | t-test | 90.0 ± 1.3 | 18 / 14 |
| AP half-width (ms) | 0.58 ± 0.01 | 19 / 11 |  | 0.66 ± 0.01 | 17 / 10 |  | 8.49E-05 | t-test | 0.64 ± 0.01 | 18 / 14 |
| Max rate of rise (mV*ms) | 521.1 ± 15.4 | 19 / 11 |  | 569.6 ± 24.7 | 17 / 10 |  | 0.09691 | t-test | 558.8 ± 22.9 | 18 / 14 |
| Max rate of decay (mV*ms) | -146.1 ± 5.7 | 19 / 11 |  | -129.5 ± 4.3 | 17 / 10 |  | 0.03942 | MW U-test | -137.1 ± 3.4 | 18 / 14 |
| 1st frequency F/I slope (Hz/pA) | 0.46 ± 0.03 | 15 / 9 |  | 0.46 ± 0.02 | 16 / 11 |  | 0.81829 | t-test | 0.44 ± 0.03 | 9 / 8 |
| Max frequency F/I slope (Hz/pA) | 0.43 ± 0.03 | 15 / 9 |  | 0.46 ± 0.03 | 16 / 11 |  | 0.53512 | t-test | 0.44 ± 0.03 | 9 / 8 |
| Spike frequency adaptation ratio (2nd/1st) | 0.82 ± 0.02 | 13 / 7 |  | 0.73 ± 0.01 | 16 / 11 |  | 0.00129 | t-test | 0.72 ± 0.02 | 9 / 8 |
| Spike frequency adaptation ratio (last/1st) | 0.64 ± 0.03 | 13 / 7 |  | 0.54 ± 0.02 | 16 / 11 |  | 0.01322 | MW U-test | 0.52 ± 0.02 | 9 / 8 |

\* See methods for an explanation of how the electrophysiological parameters were defined/measured. Data shown as mean ± s.e.m.
\* All membrane potentials were corrected for a -14 mV liquid junction potential.
\* MW = Mann-Whitney U-test; t-test = Two-sample t-test

To determine if a potential synaptic mechanism might also explain the differences in vM1 responses, we next measured both monosynaptic excitatory and disynaptic inhibitory postsynaptic currents (EPSCs and IPSCs, respectively) evoked by vM1 optical stimuli in L6a_U_ VPm-only and L6a_L_ Dual CT cell pairs recorded simultaneously or sequentially in voltage-clamp (Fig. 6*A*). Indeed, L6 contains a large diversity of GABAergic inhibitory neurons that vM1 could also engage leading to feedforward suppression of L6 CT cells (Lund et al., 1988; West et al., 2006; Kumar and Ohana, 2008; Perrenoud et al., 2012; Bortone et al., 2014; Frandolig et al., 2019). To isolate responses, EPSCs and IPSCs were recorded near the reversal potentials for inhibition and excitation, respectively (Crandall et al., 2015). For all pairs, vM1 stimulation elicited an EPSC after a brief delay (2.1 ± 0.1 ms (*n* = 28 cells, 14 pair, 9 mice), followed 1.1 ± 0.1 ms by an IPSC (Fig. 6*A*). This brief delay between EPSCs and IPSCs is consistent with monosynaptic vM1 excitation followed by inhibition elicited in a feedforward, disynaptic manner (Gabernet et al., 2005; Cruikshank et al., 2007; Crandall et al., 2015). In nearly every pair recorded (13 of 14 pairs), the EPSC was larger in the Dual CT cell than in the VPm-only CT neuron, with the average EPSC peak in Dual CT cells being ∼2.5x larger (Fig. 6*B*). We found that IPSC peaks were also larger in Dual CT cells than in VPm-only neurons (12 of 14 pairs), with the mean disyanptic IPSC of Dual CT cells being ∼2x larger (Fig. 6*C*). Both synaptic excitatory and inhibitory charges were also consistently larger in Dual CT than in VPm-only cells (Dual EPSC charge: 2.1 ± 0.3 pC; VPm-only EPSC charge: 0.8 ± 0.1 pC, n = 14 pairs, 9 mice, *p* = 0.00137, Wilcoxon paired sign-ranked test; Dual IPSC charge: 10.2 ± 1.5 pC; VPm-only IPSC charge: 4.1 ± 1.2 pC, n = 14 pairs, 9 mice, *p* = 0.00574, Wilcoxon paired sign-ranked test). Consequently, the contribution of the EPSCs to the total synaptic current/charge elicited by vM1 input (EPSC / (EPSC + IPSC)) was similar across classes of CT neurons (Dual peaks: 0.27 ± 0.04; VPm-only peaks: 0.24 ± 0.04; n = 14 pairs, 9 mice, *p* = 0.57208, Wilcoxon paired sign-ranked test; Dual charge: 0.21 ± 0.04; VPm-only charge: 0.24 ± 0.04; n = 14 pairs, 9 mice, *p* = 0.80, Wilcoxon paired sign-ranked test). Thus, these data indicate that the greater responsiveness of L6a_L_ Dual CT neurons to vM1 input results from their distinctive intrinsic properties and the strength of vM1 excitation.

**Figure 6.**
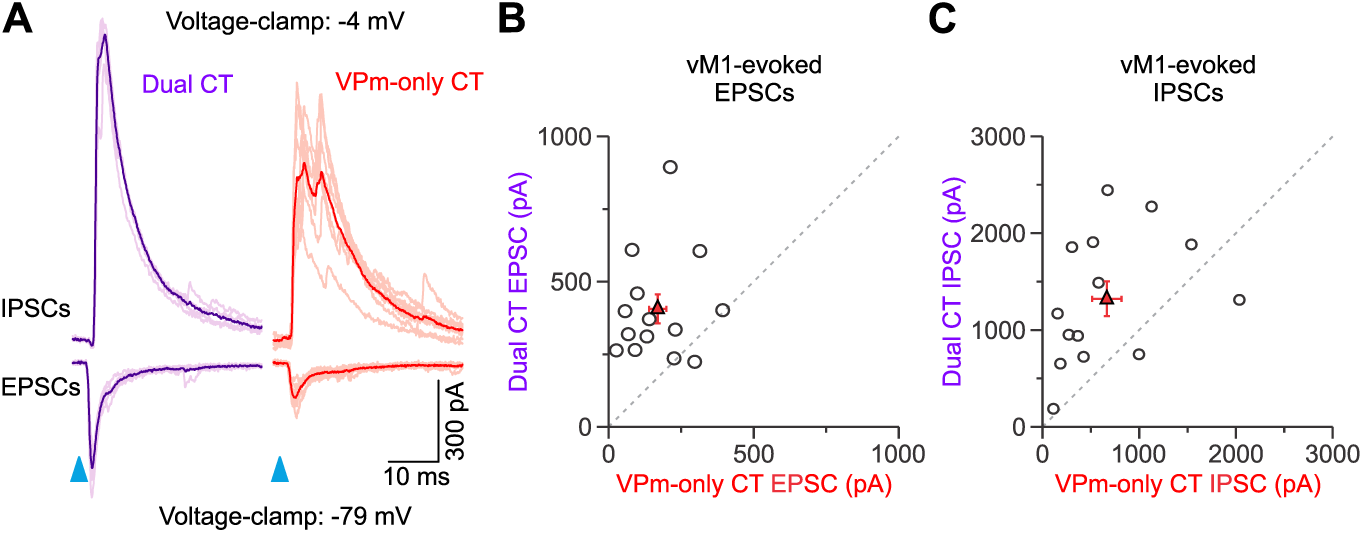
Synaptic excitation and inhibition evoked by vM1 stimulation are larger in Dual CT neurons. ***A,*** Left, Example EPSCs and IPSCs evoked by vM1 stimulation for a L6a_L_ Dual and L6a_U_ VPm-only CT cell pair. The average (dark color) and individual trials are shown (light color). EPSPs and IPSCs were recorded at the reversal potential for inhibition and excitation, respectively. ***B,*** Comparison of the EPSC peak for all pairs tested at 4x the Dual EPSP threshold (VPm-only CT: 169.4 ± 29.7 pA; Dual CT: 405.9 ± 49.6 pA; *p* = 0.0021, Wilcoxon signed-rank test; *n* = 14 pairs; 9 mice). ***C,*** Comparison of the IPSC peak for all pairs tested at 4x the Dual EPSP threshold (VPm-only CT: = 665.1 ± 152.3 pA; Dual CT: 1322.9 ± 179.6 pA; *p* = 0.00837, Wilcoxon signed-rank test; *n* = 14 pairs; 9 mice). Values are represented as mean ± SEM.

## DISCUSSION

Despite the potential power of the CT pathway, a thorough understanding of the inputs controlling CT neurons has been elusive. Here, we used the mouse vibrissal system, selective optogenetic strategies, and retrograde labeling to understand how distinct L6 CT neurons in vS1 respond to input from vM1 (Fig. 7). We found that vM1 inputs are highly selective, evoking stronger postsynaptic responses in L6a_L_ Dual VPm/POm projecting CT neurons than in L6a_U_ VPm-only projecting cells. A targeted analysis of the specific cells and synapses involved revealed that the greater responsiveness of Dual CT neurons was due to their distinctive intrinsic membrane properties and synaptic mechanisms. These data demonstrate that vS1 has at least two discrete L6 CT subcircuits distinguished by their thalamic projection patterns, intrinsic physiology, and functional connectivity with vM1. Our results also provide insights into how a distinct CT subcircuit may serve specialized roles specific to contextual modulation of tactile-related sensory signals in the somatosensory thalamus during active vibrissa movements.

**Figure 7.**
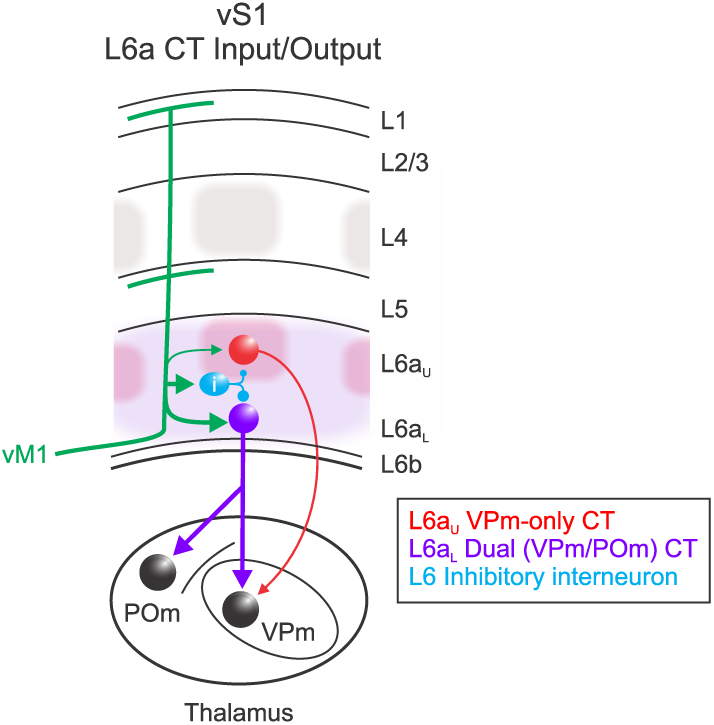
Motor modulation of distinct L6a CT circuits in vS1. VPm-only projecting CT cells are densest in L6a_U_, within infrabarrels (red shadows) that align with L4 barrels (gray) (Crandall et al., 2017). In contrast, Dual, VPm/POm, projecting CT cells are densest in L6aL (purple shadows). Dual CT cells receive strong excitatory input from vM1, whereas VPm-only CT cells receive weaker vM1 input. GABAergic inhibitory neurons, presumably in L6, are also engaged by vM1, providing disynaptic feedforward inhibition proportional to the vM1 excitation onto CT cells. Dual CT neurons are also intrinsically more excitable than VPm-only CT cells, contributing to the greater responsiveness of Dual CT neurons.

Anatomical and physiological studies in several cortical areas suggest that CT neurons are heterogeneous (Tsumoto and Suda, 1980; Fitzpatrick et al., 1994; Bourassa et al., 1995; Zhang and Deschenes, 1997; Sirota et al., 2005; Briggs and Usrey, 2009; Briggs et al., 2016; Stoelzel et al., 2017; Chevee et al., 2018; Dash et al., 2022). A prominent structural feature observed in several species and sensory systems is the separation of L6 based on CT neurons with distinct projections to the thalamus (Conley and Raczkowski, 1990; Fitzpatrick et al., 1994; Bourassa and Deschenes, 1995; Bourassa et al., 1995; Killackey and Sherman, 2003; Kim et al., 2014). This common organizational scheme suggests multiple L6 feedback pathways, each projecting to unique relay cell types. For instance, in highly visual mammals, CT circuits appear organized into separate but parallel processing streams similar to feedforward streams (Briggs and Usrey, 2009; Briggs et al., 2016).

In rodent vS1, most CT neurons in upper L6 project to VPm, the core vibrissal somatosensory nucleus, whereas the majority in lower L6 are thought to target both VPm and POm, a higher-order nucleus connected with multiple cortical and subcortical areas (Chmielowska et al., 1989; Bourassa et al., 1995; Killackey and Sherman, 2003; Chevee et al., 2018). Despite evidence for two CT classes, the proportion of each has been difficult to establish. Here, we used a double retrograde labeling strategy to quantify the proportion of each. We found that most (98%) L6a_U_ CT cells project to VPm, with 78% classified as VPm-only projecting, whereas 86% of L6a_L_ CT cells target POm, with 72% being Dual projecting. We also found that 13% of L6a_L_ CT cells are POm-only projecting. Whether these cells represent another subclass is unclear since the possibility of mislabeling exists. However, Bourassa et al. (1995) also reported cells arborizing exclusively in the POm, supporting the argument that some CT cells are POm-only projecting. These cell types, each with distinct thalamic connectivity, support the idea that vS1 has multiple CT pathways, each potentially conveying different information to targets in the thalamus. In future studies, it will be important to perform physiological recordings from each class in awake, alert subjects to determine if they have distinct response properties consistent with recent observations (Dash et al., 2022). It would also be interesting to investigate whether there are additional, morphologically unique cell types within these CT classes that could indicate further circuit-specific processing of somatosensory information (Briggs et al., 2016).

A central finding of this study is that the response to vM1 input was substantially stronger in L6a_L_ Dual CT neurons compared to L6a_U_ VPm-only cells. One mechanism contributing to the stronger response in Dual CT cells was their intrinsic properties. Although we found no difference in membrane potential, Dual CT cells were more intrinsically excitable than VPm-only cells due to their high input resistances and low rheobase. Furthermore, their lower spike threshold tended to increase their overall excitability. Another mechanism contributing to the greater responses in Dual CT cells was their stronger vM1-evoked EPSCs. On average, EPSC amplitudes in Dual CT neurons were 2.5-fold greater than in VPm-only cells. The mechanisms for the larger EPSC are unclear, but there are two possibilities: 1) the unitary synaptic connections from vM1 to Dual CT could be stronger, and or 2) more vM1 neurons might connect with each Dual CT neuron. Consistent with the innervation difference, recent work using rabies-based tracing indicates that L6a_L_ CT cells in vS1 have more presynaptic partners in vM1 than L6a_U_ CT neurons (Whilden et al., 2021). Although Dual CT cells had larger EPSCs, they also received larger IPSCs. As a result, CT cells received inhibition proportional to their vM1 excitation (Xue et al., 2014), suggesting that feedforward inhibition may contribute less to the differences in responses. Nevertheless, identifying the interneurons mediating motor integration in L6 is necessary for understanding how vM1 ultimately shapes CT feedback. These results reveal the cellular and synaptic mechanisms by which vM1 influences the excitability L6 CT neurons.

Surprisingly, the incidence of vM1-evoked spiking observed was low. However, when depolarized, responses were often large enough to trigger APs in Dual CT neurons but not VPm-only cells. How could vM1 influence CT output? One possibility is that Dual CT neurons require interactions among vM1 synapses and other excitatory inputs. For example, vM1 signals and vibrissa-sensory input could sum during active sensing, similar to vestibular and visual inputs in L6 of the visual cortex (Velez-Fort et al., 2018). Alternatively, vM1 afferents could switch CT cells to a depolarized (active) state (Zagha et al., 2013) so that sensory input is more likely to drive APs in CT neurons (Lee et al., 2008). Indeed, VPm input is relatively weak to L6 CT neurons (Crandall et al., 2017), so vM1 facilitating CT sensory responsiveness seems plausible. However, VPm responses in L6a_L_ CT cells are even weaker than those in L6a_U_ (Frandolig et al., 2019), suggesting VPm axons avoid Dual cells. Another possibility is that CT neurons integrate motor and arousal signals. There is clear evidence from studies in various sensory systems and species that arousal/state signals are present in CT neurons (Swadlow and Weyand, 1987; Steriade, 2001; Stoelzel et al., 2017; Augustinaite and Kuhn, 2020; Clayton et al., 2021; Dash et al., 2022; Reinhold et al., 2023), but direct evidence for whisker motor and arousal signal integration is lacking.

Although CT feedback circuits are ubiquitous across mammalian species and modalities (Hasse and Briggs, 2017a; Shepherd and Yamawaki, 2021), a thorough understanding of CT functions remains elusive. Indeed, previous studies suggest several functions (Contreras et al., 1996; Temereanca and Simons, 2004; Wang et al., 2006; Andolina et al., 2007; Li and Ebner, 2007; Olsen et al., 2012; Mease et al., 2014; Denman and Contreras, 2015; Hasse and Briggs, 2017b; Pauzin and Krieger, 2018; Wang et al., 2018; Pauzin et al., 2019; Ansorge et al., 2020; Born et al., 2021; Ziegler et al., 2023). This lack of understanding is likely due to complex CT circuit interactions, the nature of which is only beginning to emerge. For instance, L6 CT cells monosynaptically excite thalamic relay cells in both core sensory and higher-order nuclei, such as the VPm and POm (Scharfman et al., 1990; Golshani et al., 2001; Landisman and Connors, 2007; Lam and Sherman, 2010; Hoerder-Suabedissen et al., 2018), but CT cells also indirectly inhibit them by exciting GABAergic neurons in the thalamic reticular nucleus (TRN) (Cox et al., 1997; Kim et al., 1997; Golshani et al., 2001; Crandall et al., 2015). Evidence also suggests that the magnitude and sign of influence (enhancement or suppression) likely reflect complex interactions involving the spatial specificity and temporal properties of feedback (von Krosigk et al., 1999; Temereanca and Simons, 2004; Li and Ebner, 2007; Crandall et al., 2015; Guo et al., 2017; Hasse and Briggs, 2017b; Kirchgessner et al., 2020; Born et al., 2021; Dimwamwa et al., 2024). Such an organization could allow the cortex to influence its own sensory input in several ways. However, an open and unresolved question is whether subclasses of CT cells serve specific modulatory functions related to their organization. This question is of particular interest given the segregation in connectivity indicates that CT subcircuits are influenced by different contextual and sensory-related information.

In rodent vS1, L6a_U_ CT cells appear to project to topographically aligned VPm regions (barreloids), whereas L6a_L_ neurons have more complicated projections that include both aligned and non-aligned VPm regions as well as POm (for review, see (Deschenes et al., 1998)). Furthermore, recent work by Whilden et al. (2021) found projection differences in the TRN, with L6a_U_ and L6a_L_ CT cells targeting the core and inner edge, respectively. These TRN regions contain genetically and physiologically distinct cells that connect independently with the VPm and POm (Li et al., 2020; Martinez-Garcia et al., 2020). These studies suggest complex circuit interactions between CT subclasses and cells of the TRN and relay nuclei exist. Ultimately, to reveal an integrated dynamic picture of top-down CT influences on thalamic excitability, studying both CT circuits to understand how they operate independently and together will be critical.

## Author contributions

S.R.C. and L.E.M. designed research; L.E.M. performed the surgeries and electrophysiological experiments; D.M.A. performed the histological experiments; D.M.A. and L.E.M. performed the confocal imaging; L.E.M. and S.R.C analyzed data; L.E.M. wrote the first draft; S.R.C. wrote the final paper.

## Acknowledgments

This work was supported by NIH grants R01NS117636 (to S.R.C), R00NS096108 (to S.R.C), R00NS096108-S1 (to S.R.C), and F31NS124244 (to L.E.M.). We thank Barry Connors (Brown University) for providing the *Ntsr1-Cre* mice. Charles Lee Cox (Michigan State University) for epifluorescence imaging support. Melinda Frame and the Michigan State University Center for Advanced Microscopy for confocal imaging support.

